# Cross-species DNMT2-mediated DNA methylation with an S-phase 5mC pulse

**DOI:** 10.1101/2025.09.01.672941

**Authors:** Samia Miled

## Abstract

**Background:** The budding yeast *Saccharomyces cerevisiae* lacks endogenous DNA methyltransferases, providing a DNMT-free chassis to test catalytic potential of candidate enzymes. Building on our discovery that the *Schizosaccharomyces pombe* Dnmt2 ortholog Pmt1 can methylate DNA in vivo, we asked whether Dnmt2-family enzymes install genomic 5-methyl-2′-deoxycytidine (5mdC) across heterologous contexts.

**Results:** We fused Pmt1 to an N-terminal SNAP tag and expressed it in *S. cerevisiae*, where it was well expressed, nuclear-enriched, and non-toxic. LC/MS of genomic nucleosides revealed readily detectable 5mdC in Pmt1–SNAP cells but not detected in empty-vector controls. Following G1 arrest–release, 5mdC rose at S phase and, unlike in *S. pombe*, failed to decay over the assayed window, indicating limited methyl-cytosine clearance in the DNMT-free chassis. Extending this paradigm, expression of *Drosophila melanogaster* Dnmt2 (Dnmt2^Dm^) or *Plasmodium falciparum* TRDMT1 (TRDMT1^Pf^) in *S. cerevisiae* similarly yielded genomic 5mdC. In vivo, Drosophila manipulations supported a DNA-directed function for Dnmt2, and in vitro S2-cell assays using two independent synchronizations showed coordinated Dnmt2 transcript induction (RT-qPCR) with an accompanying methyl-DNA signal.

**Conclusion:** Across yeast, fly, and parasite orthologs, Dnmt2-family enzymes act as bona fide DNA cytosine methyltransferases. The *S. cerevisiae* chassis reveals an S-phase–linked 5mdC pulse that persists in the absence of native turnover pathways, offering a minimal, genetically tractable system to dissect substrate selection, cell-state gating, and methyl-cytosine clearance for this atypical DNMT family.

Extending this paradigm, expression of Drosophila Dnmt2 or *Plasmodium falciparum* TRDMT1 in *S. cerevisiae* similarly yielded genomic 5mdC, consistent with the first in vivo report that TRDMT1 can methylate DNA in the parasite.

## Introduction

Cytosine-5 DNA methylation (5mC) is widespread across eukaryotes; it is absent from *Saccharomyces cerevisiae*, whereas *Schizosaccharomyces pombe* was long reported to lack genomic 5mC [1–6] but is now known to harbor low-stoichiometry, Pmt1-dependent genomic 5mC detectable during S-phase re-entry from quiescence [57]. This absence in budding yeast has turned it into a powerful chassis to reconstitute and functionally probe heterologous methylation systems in a background free of endogenous cytosine methylation and its turnover pathways [5,6]. At the same time, the evolutionary history and biochemical plasticity of the DNMT2 family (also known as TRDMT1) remain unresolved: initially annotated as DNA methyltransferases by homology, DNMT2 enzymes are now established as RNA methyltransferases that catalyze C38 m5C in specific tRNAs, with roles in translation fidelity and stress tolerance across phylogeny [7–15]. In Plasmodium falciparum, TRDMT1-mediated C38 tRNA methylation supports parasite stress tolerance [58], and, critically, TRDMT1 was recently shown for the first time to methylate DNA in vivo, revealing a DNMT2-dependent DNA cytosine landscape that is modulated by oxygen availability [59]. Structural and biochemical work nevertheless indicates that the DNMT2 catalytic pocket and DNA-binding surfaces are compatible with deoxycytidine substrates, and that DNA methylation can occur under certain conditions in vitro or in cells, suggesting latent or context-dependent DNA activity [16–19].

These uncertainties intersect with long-standing debates about the existence and dynamics of cytosine methylation in insects. While multiple population-scale and mass-spectrometry studies concluded that global 5mC levels in Drosophila are extremely low or undetectable [1,20], others reported stage- or tissue-restricted signals and Tet-dependent cytosine-oxidation pathways operating in neural development and mRNA regulation [21–24]. Such discrepancies likely reflect both biology and measurement constraints: sodium bisulfite sequencing can distort estimates at very low methylation, particularly in GC-rich and damage-prone contexts [25], dot-blot/antibody approaches vary widely in specificity and dynamic range [26,27], and enzymatic methyl-seq provides improved oxidative-damage control but still requires careful calibration against orthogonal quantification [28]. In contrast, liquid-chromatography methods that directly quantify 5-methyl-2′-deoxycytidine (5mdC) relative to dC are considered robust for establishing global levels down to the 0.01–0.1% range when assay performance is validated with standards and linearity testing [29–33].

Building on our recent demonstration that the *S. pombe* DNMT2 ortholog Pmt1 can methylate DNA in vivo in its native host during S phase, we sought to test whether DNMT2 enzymes can methylate DNA when expressed in a eukaryotic genome that is otherwise devoid of cytosine methylation. We therefore established *S. cerevisiae* strains expressing N-terminal SNAP-tagged Pmt1 (Pmt1–SNAP) under conditions that preserve viability and growth, allowing precise spatiotemporal tracking of the recombinant enzyme [34–37]. We then performed cell-cycle–resolved analyses after α-factor synchronization and monitored global 5mdC accumulation by LC/MS, comparing Pmt1–SNAP with empty-vector controls. To assess generality, we reproduced the approach with *Drosophila melanogaster* Dnmt2 (Dnmt2^Dm^–SNAP) and *Plasmodium falciparum* TRDMT1 (TRDMT1^Pf^–SNAP), and complemented the heterologous yeast system with in vivo and cell-culture assays in fly embryos and S2 cells to probe stage-specific methylation and Dnmt2 expression dynamics during early development [38–48]. This two-pronged strategy leverages (i) a methylation-naïve chassis that minimizes background and turnover and (ii) orthogonal quantification by LC/MS supported by antibody-based dot blots, together enabling a stringent test of DNMT2-mediated DNA methylation across species.

Key empirical anchors for interpreting these experiments include the well-characterized timing of the early Drosophila embryonic cell cycles (rapid S/M oscillations through the syncytial blastoderm followed by lengthening cycles and cellularization) [49–52] and the known challenges of synchronizing S2 cells, which respond to temperature shifts, nutrient cues, or replication blocks with varying efficacy depending on the sub-line and protocol [41,42,53–56]. Together, these considerations motivated us to combine in vivo stages with two orthogonal enrichment regimens for S2 cells and to prioritize LC/MS for quantitative calls, using dot-blot signals for concordant trends.

## Materials and Methods

### Drosophila strains and husbandry

Three Drosophila Dnmt2 lines were used: wild-type (Dnmt2^+/+^), a *dnmt2* mutant (null), and a Dnmt2^UAS-Dnmt2^ overexpression line; all were provided by Dr. Matthias R. Schaefer. Standard husbandry was used. For overexpression, UAS-Dnmt2 was crossed to a ubiquitous GAL4 driver as indicated in the figure legends. Embryos were collected in defined windows (e.g., 0–2 h to 12–24 h), dechorionated, fixed, and stained with anti-5mC or anti-5hmC plus DNA counterstain. Imaging settings were held constant across genotypes/stages.

### Plasmid construction

The SNAP-tag coding sequence was PCR-amplified from pAct–SNAP–3×GGGs–GW (Anne Plessis lab). The SNAP amplicon was fused in-frame to the Dnmt2 open reading frame (N-terminal SNAP; linker as in the source plasmid). The ∼2.713 kb Dnmt2–SNAP fusion was excised and ligated into pRS416 (CEN/ARS, URA3) via KpnI/XhoI to yield a low-copy yeast expression vector. Analogous N-terminal SNAP fusions were generated for Dnmt2^Dm^ and TRDMT1^Pf^.

### Yeast strains, transformation, and selection

*Saccharomyces cerevisiae* wild-type strains were transformed by LiAc/PEG and selected on SC-URA plates (pRS416). Non-transformed parental strains were processed in parallel as controls. Plasmid maintenance was checked by colony PCR and growth on URA-selective media.

### SNAP labeling and fluorescence microscopy (correct timing)

Live yeast expressing SNAP fusions were labeled with 1–2 μM SNAP-Cell dye for 30 min at 30 °C, washed 3× in dye-free medium, then chased 30 min in dye-free medium before imaging. Nuclei were counterstained with Hoechst. Imaging channels: DIC, SNAP, Hoechst, and merge. Exposure and laser settings were kept constant across conditions.

### Western blotting

Whole-cell extracts were prepared in SDS sample buffer (or alkaline lysis → TCA precipitation), boiled 5 min, resolved by SDS–PAGE (10–12%), and transferred to PVDF. Membranes were blocked in 5% milk/TBST (1 h, RT) and probed overnight at 4 °C with primary antibodies as indicated: anti-Pmt1 (gift from Dr. Matthias R. Schaefer; used for Pmt1 detection) and/or anti-SNAP (for SNAP-fusion detection). Typical dilutions: 1:1,000 for primary, 1:5,000–1:10,000 HRP-conjugated secondary. Blots were developed by ECL.

### G1 synchronization and release (yeast)

Cultures were synchronized in G1 with α-factor, washed, and released into fresh medium. Samples were harvested at G1 and 30–180 min after release. DNA-content profiles (propidium iodide) were acquired on a benchtop cytometer to annotate S-phase entry.

### Genomic DNA extraction, nucleoside digestion, and LC/MS

Genomic DNA was purified by silica columns or phenol-free methods with sequential RNase treatments. Defined DNA masses were digested to nucleosides with DNase I → nuclease P1 → alkaline phosphatase and filtered prior to analysis. Reversed-phase HPLC coupled to ESI^(+)^-MS was used to quantify 2′-deoxycytidine (dC) and 5-methyl-2′-deoxycytidine (5mdC). 5mdC identity was called by co-elution with an authentic standard and diagnostic ions ([M+H]^+^ m/z 242.1; (2M+H)^+^ ∼484.3). External calibration converted areas to amounts; values are reported as %5mdC/(5mdC+5dC). ND denotes below the assay detection limit.

### Drosophila embryos: genomic DNA

preparation Staged embryos (typically 50–200 mg per prep) were collected, dechorionated in 50% household bleach for 2 min, and rinsed thoroughly in ice-cold PBS. Pellets were homogenized in lysis buffer (10 mM Tris-HCl pH 8.0, 100 mM NaCl, 25 mM EDTA, 0.5% SDS) supplemented with proteinase K (final 100–200 μg/mL) and incubated at 55 °C for 1–3 h with occasional mixing until complete lysis. The lysate was extracted twice with an equal volume of phenol: chloroform: isoamyl alcohol (25:24:1) by gentle inversion (2–3 min) followed by centrifugation (12,000 × g, 10 min, room temperature), and the upper aqueous phase was transferred each time. The pooled aqueous phase was cleaned once with an equal volume of chloroform, mixed, and centrifuged as above. Residual RNA was removed by RNase A treatment (final 100 μg/mL, 37 °C, 30 min), followed by a final chloroform extraction. DNA was precipitated by adding 0.1 volume 3 M sodium acetate (pH 5.2) and 2.5 volumes of cold 100% ethanol (or 0.7 volume isopropanol), mixed gently, and incubated at −20 °C for 30–60 min. DNA was pelleted (12,000 × g, 15 min, 4 °C), washed with 70% ethanol, air-dried briefly, and resuspended in TE (10 mM Tris-HCl pH 8.0, 0.1–1 mM EDTA) or nuclease-free water overnight at 4 °C. Yield and purity were assessed by A260/280 and A260/230 ratios; integrity was checked on agarose gels as needed. Defined masses of DNA were then carried forward to nucleoside digestion (DNase I → nuclease P1 → alkaline phosphatase) and LC/MS quantification, and to dot-blot assays as described above.

### Dot-blot for 5mC/5hmC

Denatured genomic DNA was spotted on membranes, UV-crosslinked, and probed with anti-5mC or anti-5hmC antibodies. Single-stranded DNA staining served as a loading control. Signals were compared at matched inputs.

### Drosophila S2 cells: culture, synchronization, RT-qPCR

S2 cells were maintained under standard conditions. Two synchronization regimens were used: serum deprivation followed by re-addition, and temperature shift (cold enrichment followed by re-warm). Total RNA was extracted, DNase-treated, and reverse-transcribed. Dnmt2 transcripts were quantified by RT-qPCR with housekeeping normalization; primer efficiencies were checked on standard curves.

## Results

### A DNMT-free yeast chassis reveals nuclear and cytoplasmic Pmt1–SNAP expression without toxicity

To test DNA-methyltransferase activity in a methylation-naïve background, we expressed N-terminally SNAP-tagged Pmt1 in *Saccharomyces cerevisiae*, which lacks endogenous DNA methyltransferases and detectable genomic 5mC [1,5,6]. ApoTome imaging showed punctate nuclear fluorescence for Pmt1–SNAP that overlapped Hoechst, whereas empty-vector controls displayed only background SNAP signal (Fig. 1A). Cells progressed normally through a standard α-factor G1 arrest–release regimen with indistinguishable morphology at G1, +2 h, and +4 h (Fig. 1B), and growth curves from two independent clones per genotype revealed only a modest delay in exponential entry without growth arrest (Fig. 1C). Coomassie staining indicated comparable loading across time points, and anti-SNAP immunoblotting detected a single band at the expected size for Pmt1–SNAP (Fig. 1D). These data confirm correct expression/localization and the absence of overt toxicity, enabling quantitative assays of genomic 5mC in a clean chassis [34–37,53–55].

**Figure 1.**
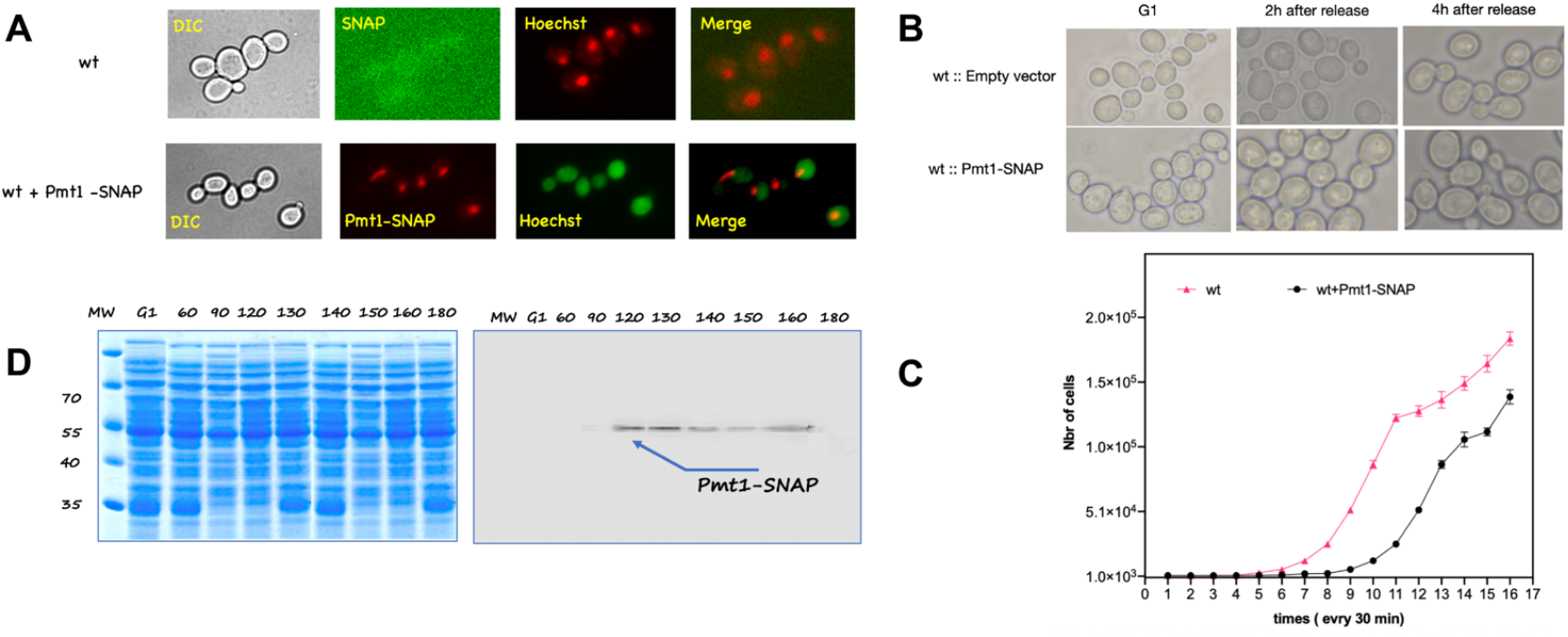
Pmt1–SNAP is expressed, nuclear-enriched, and non-toxic in *Saccharomyces cerevisiae*. **A**, ApoTome fluorescence microscopy of wt versus wt expressing N-terminally SNAP-tagged Pmt1. Channels: DIC, SNAP dye, Hoechst, and merge. Pmt1–SNAP shows punctate nuclear signal overlapping Hoechst; wt shows background SNAP staining only. **B**, Cell morphology during a G1 arrest–release experiment (G1, +2 h, +4 h after release) for wt with empty vector and wt + Pmt1–SNAP; both strains progress through the time course without abnormal morphology. **C**, Growth-curve analysis (cell numbers every 30 min). Two independent clones per genotype. Pmt1–SNAP shows a modest delay in exponential entry but no growth arrest, indicating that the construct is not toxic under the conditions used. **D**, Left, Coomassie-stained SDS–PAGE of whole-cell extracts collected along the time course (labels indicate minutes after release). Right, anti-SNAP immunoblot detects a single band at the expected size for Pmt1–SNAP across time points. Together, A–D demonstrate correct expression/localization and the absence of overt toxicity.

### An S-phase–linked rise in genomic 5mC accumulates in yeast expressing Pmt1–SNAP and does not decay over 180 min

After α-factor release, DNA-content histograms documented matched S-phase progression in wild type and Pmt1–SNAP strains (Fig. 2A), and Pmt1–SNAP remained nuclear at +120 min (Fig. 2B). LC/MS of enzymatically digested genomic DNA revealed a progressive increase in %5mdC/ (5mdC + 5dC) from G1 to 180 min in Pmt1–SNAP cells, reaching roughly 3–3.5%, while empty-vector controls remained at background (Fig. 2C). Notably, the 5mC signal did not decline within the 3-hour window, consistent with limited methyl-cytosine clearance in budding yeast, which lacks TET-type oxidases and canonical demethylation pathways [1–3,18,19]. The quantification relies on chromatographic co-elution with authentic standards and established LC/MS workflows suitable for low-stoichiometry 5mdC (ND, not detected, defined as below the assay LOD) [29–33]. Together with the S-phase timing, these data indicate that Pmt1 can methylate DNA in vivo and that the DNMT-free yeast background does not efficiently remove the modification.

**Figure 2.**
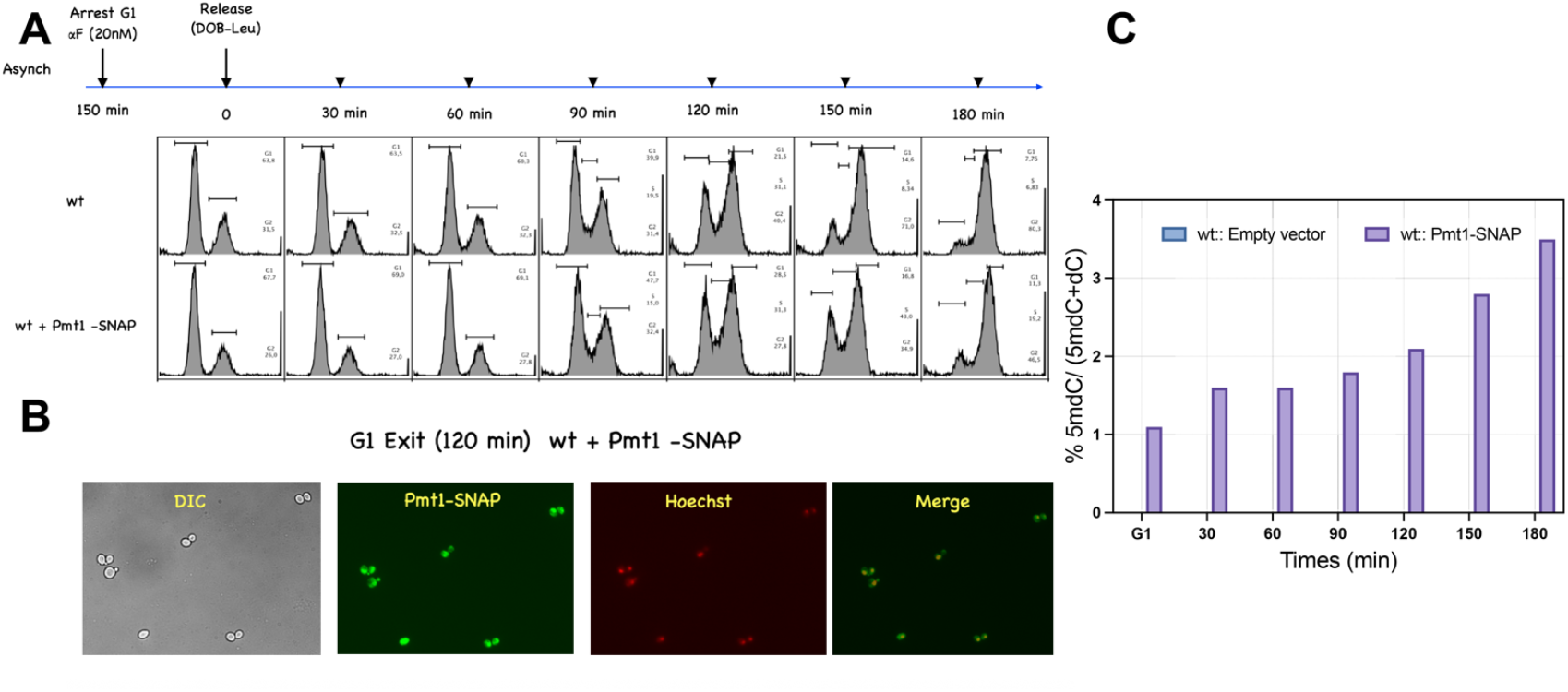
G1 arrest–release uncovers an S-phase 5mdC rise in Pmt1–SNAP cells that does not decay over 180 min. **A**, Scheme of the synchronization (α-factor G1 arrest) and release into S phase; Flow cytometry (FACS) of DNA content at the indicated times indicates equivalent cell-cycle progression in wt and Pmt1–SNAP. **B**, Microscopy at +120 min after release showing Pmt1–SNAP nuclear signal (DIC, SNAP, Hoechst, merge). **C**, LC/MS quantification of genomic 5mdC expressed as %5mdC/(5mdC+5dC) at G1 and 30–180 min after release. Empty-vector wt remains at background, whereas Pmt1–SNAP increases steadily to approximately 3–3.5% by 180 min. The absence of a downward trend indicates limited methyl-cytosine clearance in S. cerevisiae, consistent with this organism lacking endogenous DNA methylation/repair pathways for 5mdC. Bars show mean ± SD; ND indicates below assay detection.

### DNMT2 orthologs from Drosophila and Plasmodium also install genomic 5mC in the yeast chassis

To ask whether DNA-directed activity generalizes across the DNMT2 family, we expressed Drosophila Dnmt2 (Dnmt2^Dm^–SNAP) and *Plasmodium falciparum* TRDMT1 (TRDMT1^Pf^–SNAP) in the same chassis. Confocal imaging showed nuclear enrichment for both constructs (Fig. 3, top and 3A). Dot-blots of denatured genomic DNA probed with anti-5mC revealed a clear signal in transformed cells but not in non-transformed controls, with ssDNA spots confirming equal loading (Fig. 3B); we interpret dot-blots as qualitative support given known antibody variability at low 5mC [26,27]. LC/MS quantification reported increased 5mdC fractions upon heterologous expression (Fig. 3C). These results, together with the robust RNA-methyltransferase literature for DNMT2 enzymes, indicate that multiple orthologs can methylate DNA in vivo when provided chromatin access in a DNMT-free eukaryote [7–15,16–19,40,44–47].

**Figure 3.**
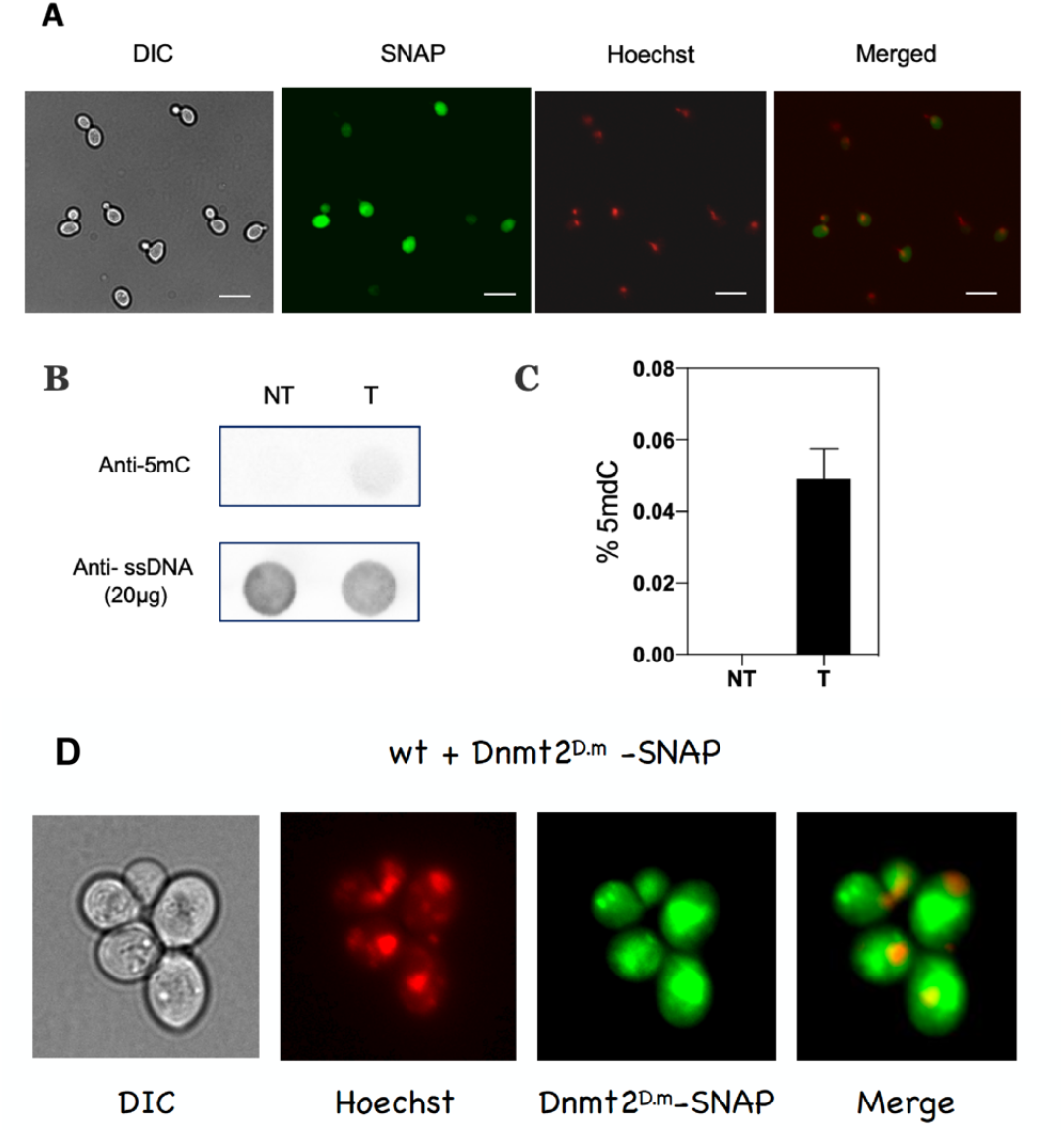
DNMT2 orthologs from Drosophila and Plasmodium install genomic 5mdC in the S. cerevisiae chassis. Top strip, Confocal images of yeast expressing Dnmt2^Dm^–SNAP (DIC, Hoechst, Dnmt2^Dm^–SNAP, merge), showing nuclear enrichment. **A**, Confocal images of yeast expressing TRDMT1^Pf^–SNAP (DNMT2^p.f^–SNAP) with the same channels; scale bar 10 μm. **B**, Dot-blot of denatured genomic DNA probed with anti-5mC, comparing non-transformed (NT) and transformed (T) yaest cells; ssDNA loading control below. Transformed cells show a clear 5mC signal. **C**, LC/MS measurement of %5mdC/ (total cytosine modifications) in NT versus T. The 5mdC fraction increases upon expression of the heterologous enzyme. *Note on nomenclature:* the figure combines Dnmt2^Dm^ (top strip) and TRDMT1^Pf^ (panels A–C); both orthologs yield detectable 5mdC in *S. cerevisiae* under the same analytical pipeline.

### Drosophila embryos show stage-dependent 5mC/5hmC signals with Dnmt2 dependence

To place the heterologous findings in organismal context, we examined early Drosophila development across the pre-blastoderm, syncytial blastoderm, cellularization, and gastrulation stages (timing summarized in Fig. 4A) [49–52]. Whole-mount immunofluorescence showed stronger 5mC signal in Dnmt2^+/+^ embryos than in the dnmt2 line presented, with merged images confirming DNA overlap (Fig. 4B). Stage-matched dot-blots probed with anti-5hmC (and in parallel with anti-5mC) revealed time-dependent signals in Dnmt2^+/+^ that were diminished or absent in dnmt2^−/−^, with ssDNA controls uniform across samples (Fig. 4C–D). Independent HPLC quantification reported a peak in 5mdC near 6–8 h (∼6%) and a transient 5hmdC rise that subsided by 12–24 h (Fig. 4E). Although global 5mC in Drosophila is near the detection floor of many assays [1,20,25], these convergent signals - together with reports of Tet-dependent cytosine-oxidation pathways in specific developmental settings - support the view that DNA cytosine modification is present at low but regulated levels and that Dnmt2 contributes to this landscape in vivo [21–24,25–27,29–33].

**Figure 4.**
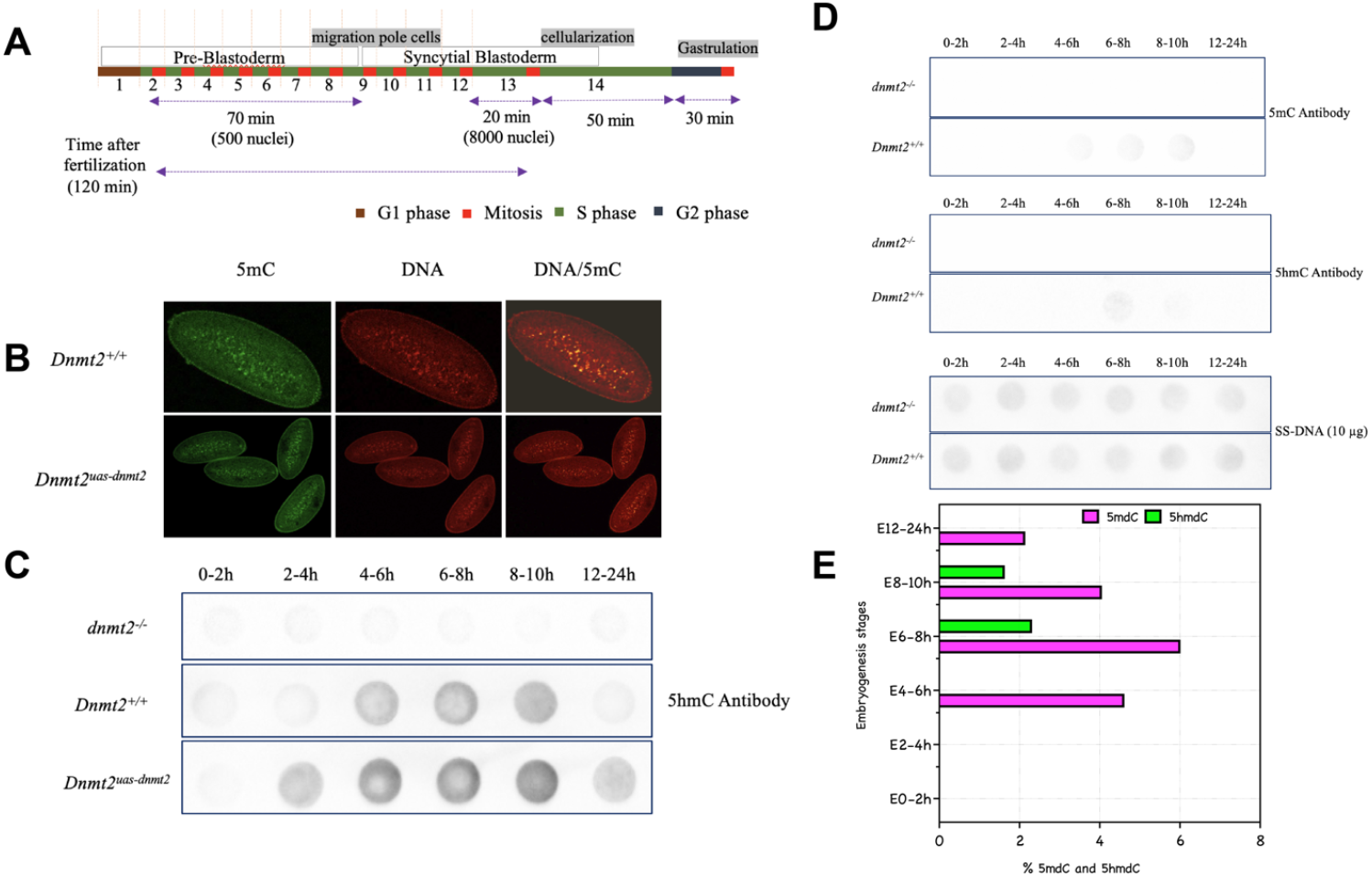
Drosophila embryos: developmentally regulated DNA 5mC/5hmC signals that depend on Dnmt2. **A**, Developmental timeline from fertilization through pre-blastoderm, syncytial blastoderm, cellularization, and gastrulation, with approximate durations and predominant cell-cycle phases. **B**, Whole-mount immunofluorescence of embryos stained for 5mC (green) and DNA (red) with merged images, comparing Dnmt2^+/+^ to the line denoted Dnmt2^uas-dnmt2^ (as labeled on panels); Dnmt2 competence correlates with stronger 5mC signal. **C**, Dot-blot across developmental windows (0–2 h to 12–24 h) probed with anti-5hmC for dnmt2^−/−^, Dnmt2^+/+^, and Dnmt2^uas-dnmt2^; intensity tracks Dnmt2 status. **D**, Independent dot-blot series probed with anti-5mC (top) and anti-5hmC (middle) in dnmt2^−/−^ and Dnmt2^+/+^, with ssDNA loading controls (bottom). Loss of Dnmt2 reduces or abolishes the immunoreactivity, while wild type retains a time-dependent signal. **E**, HPLC quantification during early embryogenesis showing a 5mdC peak around 6–8 h (∼6%) and a transient 5hmdC rise (∼2–2.5%) that declines by 12–24 h. Together, B–E support a DNA-directed Dnmt2 activity in vivo during embryogenesis.

Our heterologous TRDMT1^Pf^ data in the DNMT-free yeast chassis mechanistically extend the initial in vivo report that TRDMT1 can methylate DNA in *P. falciparum* [59] and complement its established role in tRNA-mediated stress tolerance [58].

### Two orthogonal S2-cell synchronizations link Dnmt2 induction to increased methyl-DNA signal at G1 exit

We next used two standard synchronization strategies in Drosophila S2 cells. In serum-deprivation, cells accumulated in G1 after 24 h without serum and re-entered the cycle 16 h after serum addition; flow cytometry validated the transitions (Fig. 5A, a1). RT-qPCR showed induction of Dnmt2 transcripts upon G1 exit (Fig. 5A, a2), and anti-5mC dot-blots indicated stronger methyl-DNA signal at exit than in asynchronous or G1-held cells (Fig. 5A, a3). A temperature-shift regimen (1 h, at 4 °C followed by 16 h at 22 °C) produced comparable G1 enrichment and exit by flow (Fig. 5B, b1), accompanied by similar Dnmt2 induction (Fig. 5B, b2) and a rise in 5mC by dot-blot (Fig. 5B, b3). While S2 synchronization efficiencies vary with sub-line and protocol [41–43,53–56], the convergence of two independent approaches supports coupling between Dnmt2 expression and methyl-DNA accumulation at cell-cycle re-entry.

**Figure 5.**
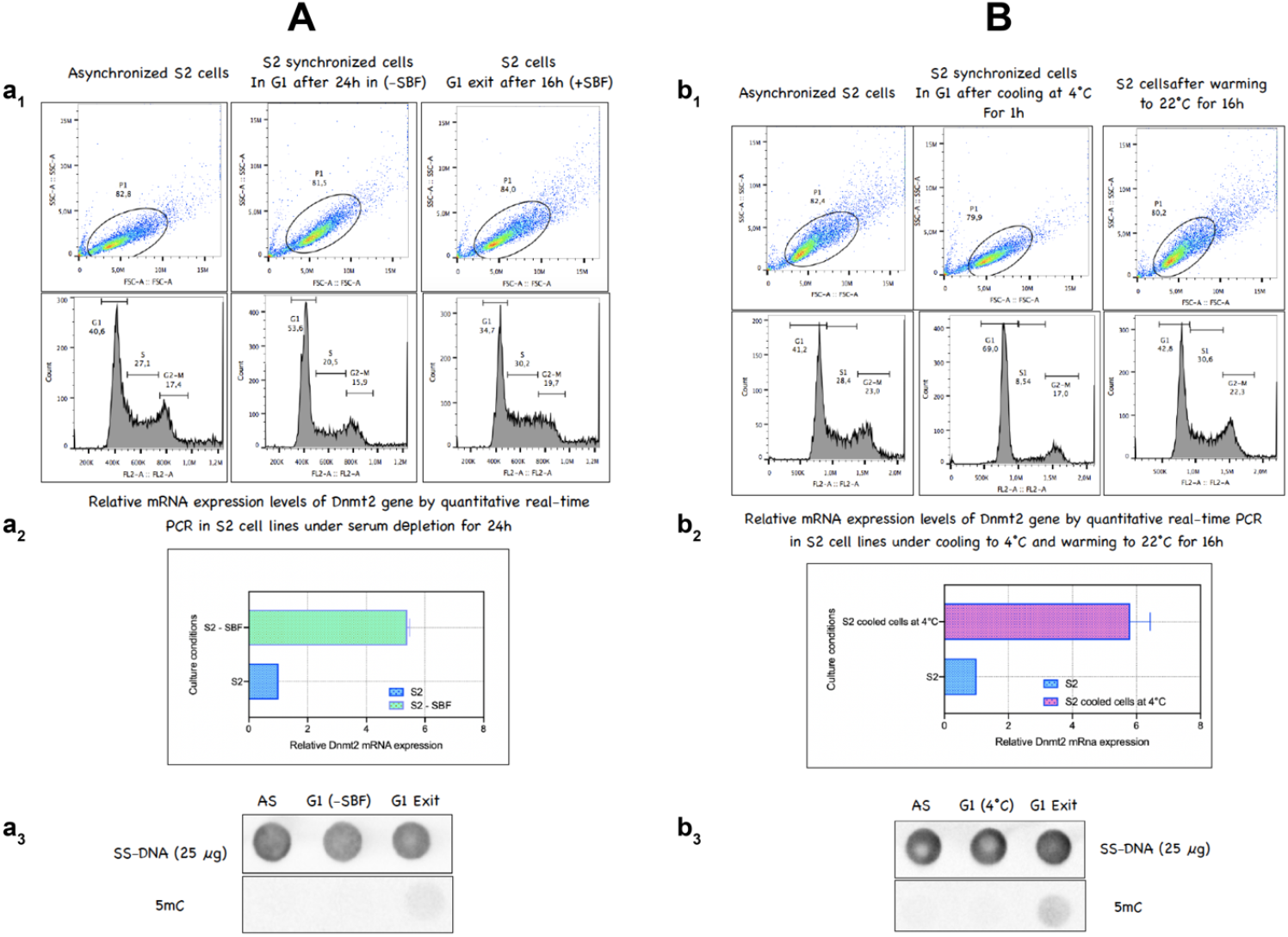
Drosophila S2 cells: two orthogonal synchronization protocols reveal Dnmt2 transcript dynamics and DNA 5mC signal. **A**, Serum-depletion protocol. **a1**, Flow cytometry of asynchronous cells (AS), cells held in G1 after 24 h in serum-free medium (−SBF), and G1 exit 16 h after serum re-addition (+SBF). **a2**, RT-qPCR quantification of Dnmt2 mRNA (normalized to housekeeping controls) across the same states; expression increases upon G1 exit. **a3**, Dot-blot of genomic DNA probed with anti-5mC (ssDNA loading above), showing higher 5mC at G1 exit compared with AS and G1(−SBF). **B**, Temperature-shift protocol. **b1**, Flow cytometry of AS cells, cells cooled at 4 °C for 1 h (G1 enrichment), and cells warmed to 22 °C for 16 h (G1 exit). **b2**, RT-qPCR of Dnmt2 mRNA across states; induction accompanies exit. **b3**, Anti-5mC dot-blot with ssDNA loading indicates increased methyl-DNA signal upon exit. The convergence of the two protocols supports a coupling between Dnmt2 expression and DNA 5mC accumulation at cell-cycle re-entry.

Across a methylation-naïve yeast chassis and native Drosophila contexts, these experiments show that DNMT2-family enzymes - including Pmt1, Dnmt2^Dm^, and TRDMT1^Pf^ - an install genomic 5mC in vivo; that the signal rises with S-phase entry and persists in budding yeast lacking demethylation pathways; and that in Drosophila embryos and S2 cells, Dnmt2-dependent cytosine modifications vary with developmental stage and re-entry from G1. The combined imaging, LC/MS, and dot-blot data provide a cross-system framework to dissect substrate selection, chromatin access, and methyl-cytosine turnover for DNMT2 enzymes [7–19,21– 24,26–33,41–47,49–56].

## Discussion

Reconstituting DNMT2 orthologs in a methylation-naïve eukaryote reveals that Pmt1 from *S. pombe*, Dnmt2^Dm^ from *Drosophila*, and TRDMT1^Pf^ from *Plasmodium* are each capable of installing detectable 5mdC in genomic DNA when expressed in *S. cerevisiae*. The chassis lacks endogenous 5mC and enzyme systems that might confound installation or removal [1–6], while in its native host *S. pombe*, Pmt1-dependent genomic 5mC has been detected at low stoichiometry during G0→S [57]. Notably, our cross-species findings dovetail with the first in vivo demonstration that TRDMT1 can methylate DNA in *P. falciparum* under variable oxygen conditions [59], complementing its established role in tRNA-mediated stress tolerance [58].

LC/MS-based quantification in the chassis, together with imaging and Drosophila assays, supports a conserved DNA-directed activity for DNMT2 enzymes. Second, we verified expression, nuclear localization and non-toxicity of SNAP-tagged constructs, and followed cell-cycle progression after α-factor release using standard budding-yeast synchronization, minimizing physiological artifacts [34–37,57–59]. Third, LC/MS quantification was anchored by calibration curves and detection-limit determinations and thus provides an orthogonal metric that is insensitive to the well-documented biases of bisulfite chemistry and the variability of antibody reagents at very low methylation levels [25–33]. The monotonic increase of 5mdC following G1 release in Pmt1–SNAP yeast argues that DNMT2-dependent DNA methylation can occur during S phase, consistent with our earlier observations in *S. pombe* and with the long-standing mechanistic expectation that DNA engagement by DNMT2 uses the same catalytic motifs that act on tRNA [13,16–19].

A second inference is that budding yeast clears DNMT2-installed 5mC poorly. In our time-course, 5mdC levels failed to return to baseline after replication, implying that removal is inefficient over the experimental window. This is compatible with the absence of TET enzymes and other cytosine-oxidation/repair pathways that mediate active demethylation in animals and plants [2,18,19]. It also suggests that any apparent decline of global 5mdC at longer times in other systems may reflect cellular turnover or dilution rather than enzymatic erasure. Future mapping by EM-seq or enzymatic mTAG-based approaches could define sequence context and replication-fork proximity of the DNMT2-dependent marks in the yeast genome [28].

Our heterologous results generalize to two others eukaryotic DNMT2s. Dnmt2^Dm^ and TRDMT1^Pf^ both yielded positive LC/MS signals and immuno-dot blots in *S. cerevisiae*, reinforcing the view that DNMT2’s substrate scope includes deoxycytidines in vivo given appropriate chromatin access. For TRDMT1^Pf^, which is firmly established as a tRNA-C38 methyltransferase with critical roles in parasite stress tolerance and development, this DNA activity aligns with structural predictions and selected in vitro reports [15,16,44–47]. These findings do not negate the dominant RNA function of DNMT2 enzymes, but they demonstrate that DNA methylation is a latent and evolvable property of the DNMT2 fold.

The Drosophila experiments provide organismal context. In embryos, stage-matched dot blots and LC/MS revealed low but dynamic 5mdC (and 5hmC) signals across the syncytial blastoderm and early cellularization, peaking during windows rich in DNA synthesis. While Drosophila global 5mC levels are near the detection floor of many methods [1,20,25], our data converge with reports that cytosine-modifying pathways - including Tet-dependent chemistry - operate in specific developmental contexts in flies, especially in the nervous system [21–24]. In S2 cells, two orthogonal enrichment regimens (nutrient withdrawal and temperature shift) increased Dnmt2 transcript levels and methylated-DNA dot-blot signals, consistent with well-known coupling of cell-cycle programs to metabolic and environmental cues in insect cells [41–43,53–56]. Given the recognized difficulty of achieving perfect synchrony in S2 sub-lines, we interpret these results as evidence that Dnmt2 expression and global cytosine methylation can co-vary with cell-cycle/physiological state, not as precise measurements of phase-specific flux.

Mechanistically, how might DNMT2 access DNA? Structural analyses indicate that the DNMT2 catalytic cysteine and cofactor-binding architecture overlap with DNA-methyltransferase chemistry; DNA binding has been observed, and DNA methylation can be promoted when tRNA competition is reduced or when chromatin exposes DNMT2-preferred motifs [16–19]. Our S-phase–linked accumulation in yeast fits a model in which transient single-stranded or under-wound DNA at replication forks or repair intermediates provides entry points for DNMT2. The persistence of 5mdC in yeast further implies that once installed, the modification is not rapidly excised. These mechanistic hypotheses can be tested by (i) mapping 5mdC at base resolution in the yeast genome under controlled replication perturbations, (ii) mutationally swapping RNA/DNA binding surfaces in DNMT2 orthologs, and (iii) competition experiments with queuosine-modified tRNAs known to regulate DNMT2 activity [14,47].

Conceptually, placing the parasite evidence for TRDMT1-driven DNA methylation [59] alongside the yeast chassis and fly data reframes DNMT2 as a context-gated writer of DNA 5mC. The dominant physiological role in tRNA methylation and stress responses [58] likely masks DNA activity in many settings; however, when chromatin is accessible, competition from tRNA is reduced, and metabolic conditions are permissive, DNMT2 can mark DNA at low stoichiometry. This conditional capacity may be biologically sufficient if deployed transiently at critical transitions (for example, G0→S or developmental windows), as seen in S. pombe [57] and suggested by the S-phase behavior in our chassis.

In sum, DNMT2-family enzymes from yeast, fly, and parasite backgrounds can methylate DNA in vivo. The *S. cerevisiae* chassis reveals an S-phase 5mC accumulation that persists in the absence of native turnover, while parasite work provides the first in vivo precedent for TRDMT1-mediated DNA methylation under physiologically relevant conditions [59]. These convergent lines of evidence motivate base-resolution mapping and mechanistic dissection of how cell state, chromatin access, and metabolic cues gate DNMT2 access to DNA.

### Data Origin, Figures, and Materials Availability

The data presented in this manuscript were generated as part of an independent research program that I conceived and conducted while serving as a Temporary Teaching and Research Assistant (ATER) at Université Paris Cité, affiliated with the Jacques Monod Institute (IJM). This preprint reports the heterologous validation of DNMT2-family DNA methyltransferase activity in a DNMT-free *Saccharomyces cerevisiae* chassis and complementary assays in Drosophila and Plasmodium systems. By depositing this work on bioRxiv, I aim to ensure permanent accessibility for the scientific community and to establish a clear, traceable record of authorship.

All figures, datasets, and interpretations are the intellectual property of the author and acknowledged collaborators. This preprint is released under the Creative Commons Attribution (CC BY 4.0) license selected at submission. Any reproduction, reuse, or derivative work - including figures, datasets, or textual excerpts - must cite this preprint using its DOI and full bibliographic reference.

## Acknowledgments

I am especially thankful to Dr. Sylvie Pochet and Frédéric Bonhomme (Unité de Chimie et Biocatalyse, Institut Pasteur) for their exceptional technical expertise in the LC/MS validation experiments. I also thank Dr. Benoît Arcangioli (Unité Dynamique du Génome, Institut Pasteur) for his scientific guidance and insightful discussions. I am deeply grateful to Prof. Anne Plessis (Institut Jacques Monod), my lab head, for her help with the Drosophila developmental work and for enabling these experiments. I thank Matthieu Sanial for assistance in establishing SNAP-tag methods, and Isabelle Becam for expert help with Drosophila larval dissections. Finally, I thank Dr. Matthias R. Schaefer for providing the Drosophila Dnmt2 construct and the anti-Pmt1 antibody.

